# Strengthening *Bordetella pertussis* genomic surveillance by direct sequencing of residual positive specimens

**DOI:** 10.1101/2023.12.08.570824

**Authors:** Yanhui Peng, Margaret M. Williams, Lingzi Xiaoli, Ashley Simon, Heather Fueston, M. Lucia Tondella, Michael R. Weigand

**Author notes:** Address correspondence to Michael R. Weigand,. MW: Division of Healthcare Quality Promotion, National Center for Emerging and Zoonotic Infectious Diseases, Centers for Disease Control and Prevention, Atlanta, GA, USA. AS: Division of Parasitic Diseases and Malaria, Global Health Center, Centers for Disease Control and Prevention, Atlanta, GA, USA. LX: Division of Foodborne, Waterborne, and Environmental Diseases, National Center for Emerging and Zoonotic Infectious Diseases, Centers for Disease Control and Prevention, Atlanta, GA, USA.

## Abstract

Whole-genome sequencing (WGS) of microbial pathogens recovered from patients with infectious disease facilitates high-resolution strain characterization and molecular epidemiology. However, increasing reliance on culture-independent methods to diagnose infectious diseases has resulted in few isolates available for WGS. Here we report a novel culture-independent approach to genome characterization of *Bordetella pertussis*, the causative agent of pertussis and a paradigm for insufficient genomic surveillance due to limited culture of clinical isolates. Sequencing libraries constructed directly from residual pertussis-positive diagnostic nasopharyngeal specimens were hybridized with biotinylated RNA “baits” targeting *B. pertussis* fragments within complex mixtures that contained high concentrations of host and microbial background DNA. Recovery of *B. pertussis* genome sequence data was evaluated with mock and pooled negative clinical specimens spiked with reducing concentrations of either purified DNA or inactivated cells. Targeted enrichment increased yield of *B. pertussis* sequencing reads up to 90% while simultaneously decreasing host reads to less than 10%. Filtered sequencing reads provided sufficient genome coverage to perform characterization via whole-genome single nucleotide polymorphisms (wgSNP) and whole-genome multilocus sequencing typing (wgMLST). Moreover, these data were concordant with sequenced isolates recovered from the same specimens such that phylogenetic reconstructions from either consistently clustered the same putatively linked cases. The optimized protocol is suitable for nasopharyngeal specimens with IS*481* Ct < 35 and > 10 ng DNA. Routine implementation of these methods could strengthen surveillance and study of pertussis resurgence by capturing additional cases with genomic characterization.

## Introduction

Whooping cough (‘pertussis’) remains a persistent public health challenge with highest rates of morbidity and mortality reported in young infants (1). Although coverage with pertussis-containing vaccines among children remains high, cases increased steadily in the United States from the late 1980s until the start of the COVID-19 pandemic in 2020. Waning protection conferred by acellular vaccines, adopted by many industrialized countries beginning in the 1990s, likely contributes to increased disease among vaccinated individuals, more so than longer-lasting protection offered by whole-cell formulations (2, 3). The causative agent, *Bordetella pertussis*, exhibits little molecular variation and recent mutations to genes encoding immunogenic proteins included in current acellular pertussis vaccines have swept the population quickly, evidence of likely vaccine-driven selection (4–6). However, additional questions regarding the role of pathogen ecology and evolution in pertussis resurgence remain unanswered (7, 8).

High-throughput sequencing has transformed public health microbiology (9) and recent whole-genome sequencing (WGS) analyses of *B. pertussis* have revealed new insights about the pathogen’s population structure and dispersion through high-resolution characterization of clinical isolates (5, 10–13). But effective pertussis genomic surveillance, and subsequent study of pathogen contributions to disease resurgence, is limited by the declining use of diagnostic culture as fewer clinical isolates are available for WGS (14). Multiple influences lead to the decrease in cultured *B. pertussis* isolates, such as adoption of culture-independent diagnostic tests (CIDTs) and multi-pathogen respiratory panels, reliance on commercial testing providers, and stability of selective transport media, some of which limit culture of microbial pathogens more broadly (15). Additionally, the specificity of different pertussis diagnostic methods varies with patient age, timing of specimen collection (i.e., in relation to cough onset), and vaccination status (16). As a result, Enhanced Pertussis Surveillance (EPS) as part of the Emerging Infections Program (EIP), which conducts systematic case ascertainment and augmented data collection across seven states that include ∼7.0% of the US population, recovers *B. pertussis* isolates for, on average, fewer than 3.5% of captured cases annually (17).

Shotgun metagenomics can identify individual microbes within complex mixtures (18) and computational techniques now allow recovery of metagenome-assembled genomes (MAGs) from environmental samples (19, 20). Although metagenomics promises a pathogen-agnostic assay platform for detection as well as characterization, clinical applications routinely face the two-fold challenge of low target abundance and high background contamination from the host patient and co-occurring microorganisms(21, 22). In practice, many samples collected for infectious disease diagnosis, including nasopharyngeal (NP) specimens for suspected pertussis, are heavily loaded with human cells or exogenous DNA, limiting recovery of microbial genomic information and thus assay sensitivity. As a result, multiple laboratory approaches have been introduced to improve microbial pathogen recovery, including selective lysis (23, 24), whole genome amplification (25–27), and target enrichment (28). Successful application of these tools has enabled molecular characterization for improved outbreak investigation and surveillance (29–31). For pertussis specifically, thus far selective lysis treatment of NP specimens has shown only modest improvement in recovery of *B. pertussis* genomic data (23). Alternatively, the relatively limited genetic diversity of *B. pertussis* may benefit from target enrichment or amplification approaches that rely on existing genomic data to synthesize RNA probes or oligo primers, provided the added cost can be minimized (28, 32).

Here we present the development of novel methodology for culture-independent whole-genome sequencing (CIWGS) of *B. pertussis* from residual, positive diagnostic nasopharyngeal specimens, reducing reliance on culture for genomic data. Target enrichment via in-solution hybridization with a custom whole-genome RNA bait library yielded sequencing reads that comprised up to 90% *B. pertussis* genomic data while simultaneously reducing host contamination to below 10%. The resulting data were sufficient for robust genomic characterization concordant with that of cultured *B. pertussis* isolates using standard bioinformatic tools. These results demonstrate that CIWGS directly from residual specimens can enable molecular characterization of more pertussis cases, improving genomic surveillance and study of *B. pertussis* contributions to disease resurgence.

## Results

### Laboratory optimization

The CIWGS laboratory workflow was designed to complement existing, validated protocols for pertussis diagnosis from NP specimens. A schematic overview of the workflow is presented in FIGURE 1 and centers around hybridization of RNA “baits” to sequencing library fragments that contain matching target *B. pertussis* genomic fragments. RNA bait libraries were prepared from a mixture of 10 *B. pertussis* isolates selected to cover the breadth of genomic diversity common among the bacterial population (Table S1). Laboratory parameters for target enrichment were optimized using pooled PCR-positive clinical specimens as input material for library preparation. Performance of RNA bait hybridization was tested at various temperatures and DNA sequencing library inputs to maximize *B. pertussis* enrichment, which was evaluated by real-time PCR assay (Figure S1).

**Figure 1.**
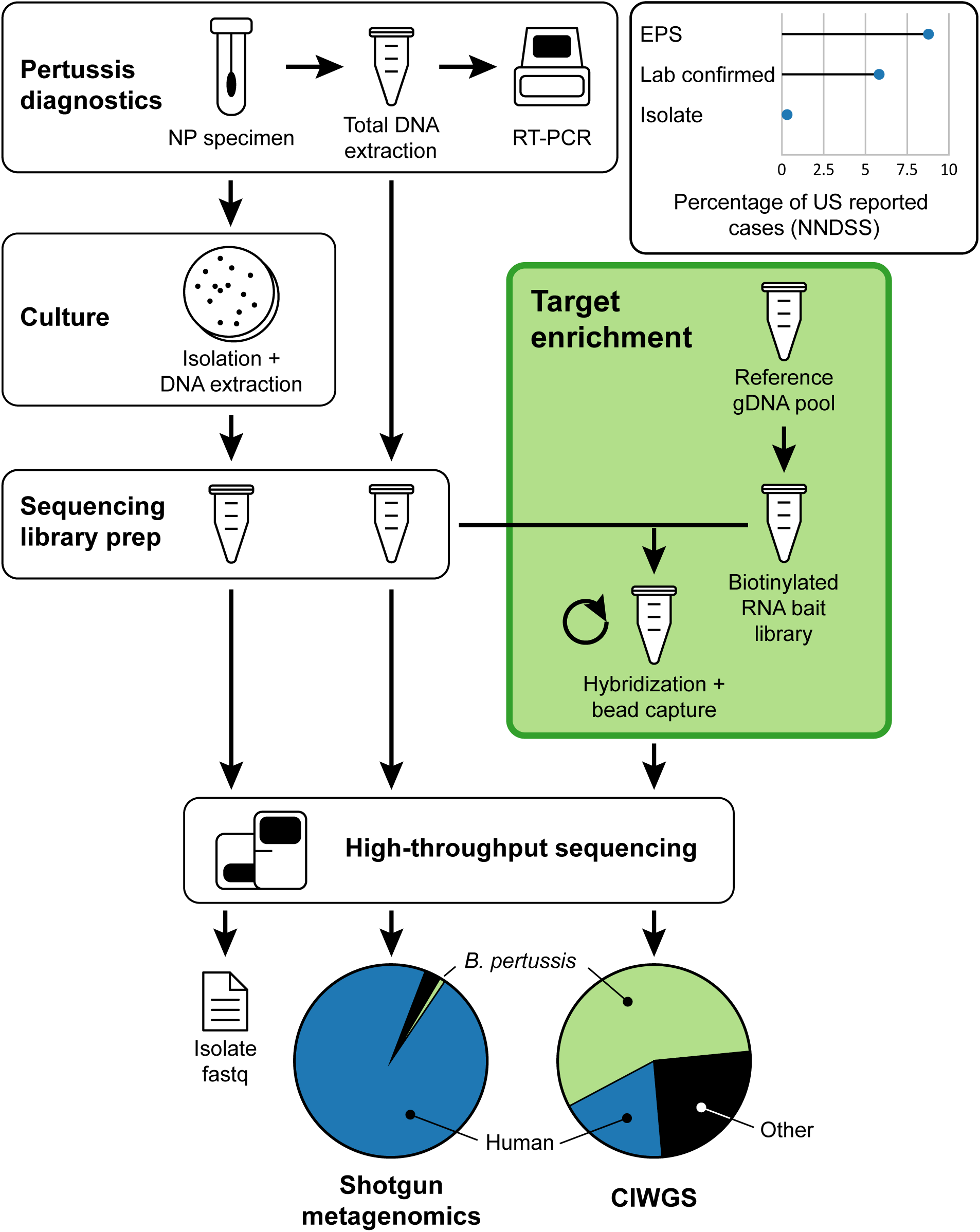
Schematic overview of the CIWGS workflow, which utilizes a RNA bait library for target enrichment of *B. pertussis* genome fragments within a sequencing library prepared directly from DNA extracts of residual nasopharyngeal (NP) species used for diagnostics. RNA baits are prepared from a diverse mixture of *B. pertussis* isolates (Table S1) and target enrichment yields significantly more sequencing reads containing *B. pertussis* genomic data than shotgun metagenomics. Less than 4% of annual reported pertussis cases are culture positive. This approach increases case coverage for genomic surveillance (inset).

Enrichment fold changes after RNA bait hybridization were calculated from *IS481* Ct values and normalized with the concentrations of the DNA libraries (Figure S1). The highest enrichment fold (3,079 x) was observed at hybridization temperature of 67.5 °C and greater DNA sequencing library input yielded more *B. pertussis* DNA fragments captured by the RNA bait library (Figure S1).

### Recovery from spiked specimens

Recovery of *B. pertussis* genome sequence data was first assessed with a series of mock specimens prepared from a mixture of commercial human DNA and a microbiome standard spiked with reducing concentrations (n=20) of extracted *B. pertussis* isolate DNA. Enrichment was evaluated by mapping sequencing reads to an existing reference-quality genome assembly for the same isolate and sequencing the background mock mixture produced only reads that mapped to regions of the rRNA operon highly conserved among bacteria. The proportion of *B. pertussis* sequencing reads was consistently enriched to an average 76% at spiked fractions down to 0.001 (IS*481* Ct = 21.1) and additional reads could be recovered at lower concentrations with a subsequent round of enrichment (FIGURE 2A). Mapping filtered *B. pertussis* sequencing reads and quantifying coverage breadth indicated that over 98% of the expected 4.1 Mbp were recovered with at least 20X depth at spiked fractions down to 0.001 and at least 90% at fraction 0.0001 (IS*481* Ct = 24.5) following a second enrichment (FIGURE 2B).

**Figure 2.**
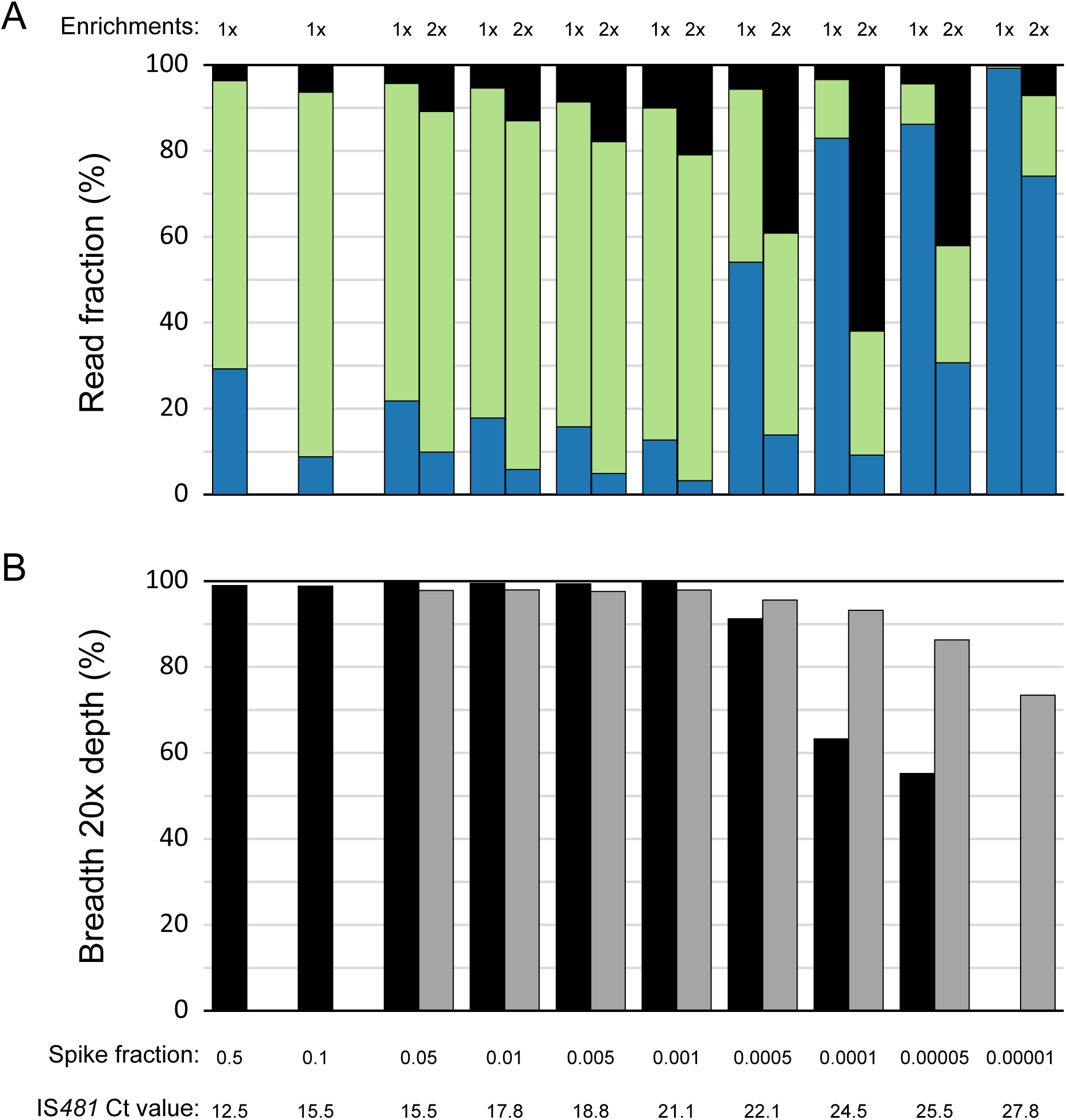
*B. pertussis* genomic data recovery from mock specimens spiked with reducing concentrations of isolate DNA following either 1x or 2x enrichments. (A) The fraction of sequencing reads containing *B. pertussis* (green), human (blue), or other (black) genomic data. (B) Coverage breadth at 20X depth across a *B. pertussis* reference genome assembly at 1x (black) or 2x (grey) enrichments.

Sequence recovery was further evaluated in a similar manner by spiking *B. pertussis* cells at reducing concentrations (n=23) into a pooled mixture of pertussis-negative NP aspirates, the background microbial composition of which was devoid of any *Bordetella* species with only spurious detection by metagenomic read classification (TABLE S2). Target enrichment yielded significant recovery of *B. pertussis* sequencing reads (Figure 3A). Average coverage breadth at 20x depth of at least 98% was observed down to spike fraction of 0.0001 (IS*481* Ct = 27.3) with 1x enrichment, and above 95% down to 0.000001 (IS*481* Ct = 35.3) with 2x enrichments (Figure 3B). Filtered *B. pertussis* reads were sufficient for downstream characterization by wgMLST, exceeding the minimum 3,000 allele call threshold established previously (Figure 3C)(33). Detailed metrics are provided in Table S3. By comparison, shotgun metagenomic (unenriched) sequencing of residual NP specimens with IS*481* Ct = 11-28 produce reads almost entirely composed of human genomic data and covered < 1% of *B. pertussis* genome nucleotides with 20X depth (Table S4). Direct comparison of residual clinical specimens sequenced in parallel with and without target enrichment further illustrated the improved *B. pertussis* genome recovery of the CIWGS workflow (Table S5).

**Figure 3.**
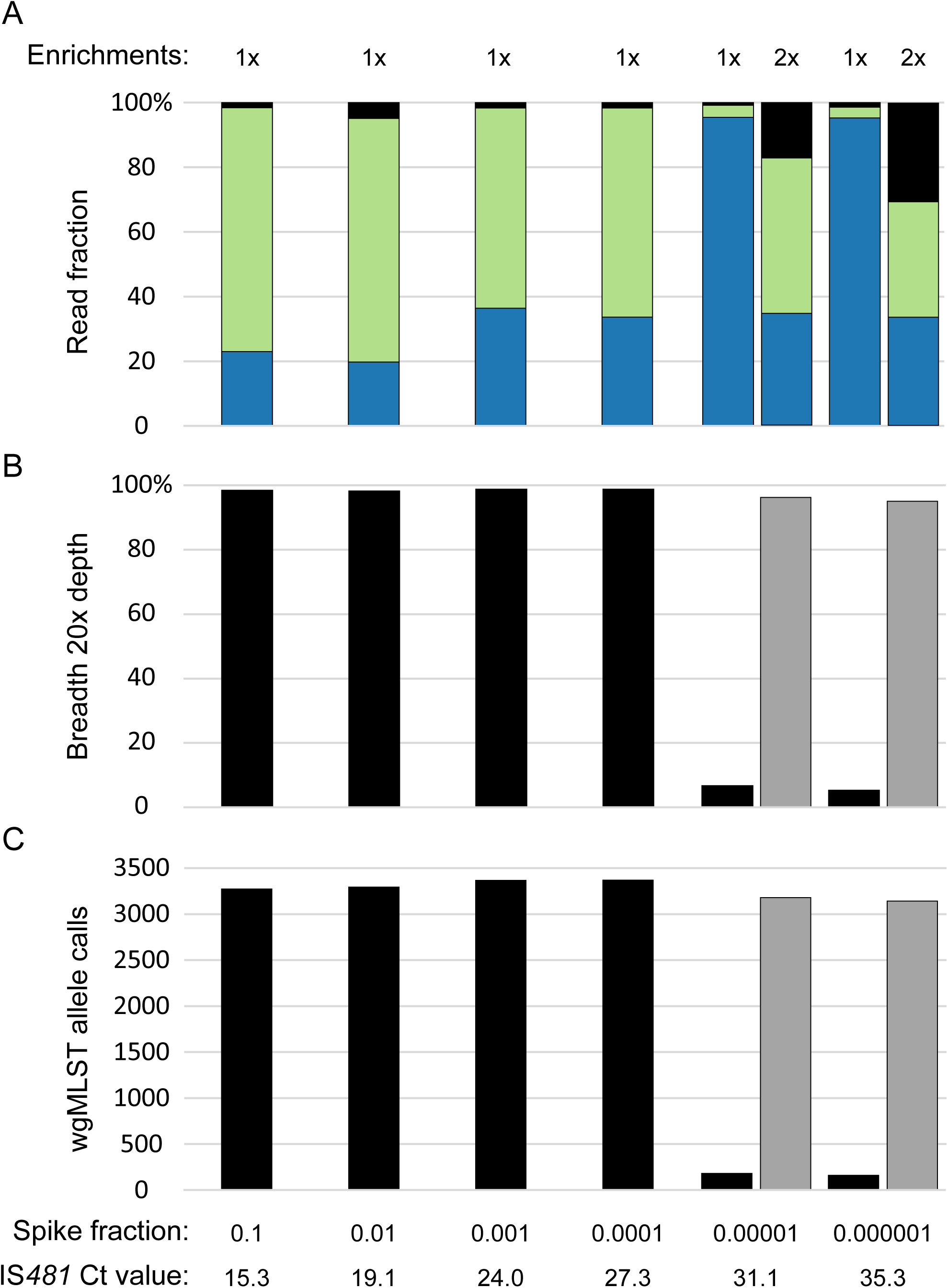
*B. pertussis* genomic data recovery from pooled negative clinical specimens spiked with dilutions of isolate cells following either 1x or 2x enrichments. (A) The fraction of sequencing reads containing *B. pertussis* (green), human (blue), or other (black) genomic data. (B) Coverage breadth at 20X depth across a *B. pertussis* reference genome assembly following 1x (black) or 2x (grey) enrichments. (C) wgMLST allele calls following 1x (black) or 2x (grey) enrichments.

### Accuracy and specificity

The accuracy of CIWGS was assessed by comparing read data from spiked specimens, both mock mixtures (n=16 positive) and pooled pertussis-negative NP aspirates (n=16 positive, n=10 negative) described above, to previous sequence data from the added *B. pertussis* isolate. Sequencing reads were first filtered by subtractive mapping to a human reference assembly and then mapped to the existing reference-quality genome assembly matching the added *B. pertussis* isolate. In all cases, 0 single nucleotide polymorphism (SNP) differences were detected when mapping filtered CIWGS reads from spiked specimens to the corresponding genome assembly. The calculated percent agreement was 100% (TP = 32/32) and the overall percent agreement was 100% (TP = 32, TN = 10, FP = 0, FN = 0), according to formulas listed in the methods. Additionally, 0 SNP differences were observed in direct comparison of CIWGS reads from 1x- and 2x-enrichment of the same spiked specimen (Table S6).

The specificity of CIWGS for *B. pertussis* was evaluated by separately spiking pooled pertussis-negative NP aspirates with extracted DNA from two related species, *B. holmesii* and *B. parapertussis*. Hybridization with the *B. pertussis* RNA baits did capture and enrich sequencing library fragments from both *B. holmesii* C690 and *B. parapertussis* J859, whose genome-wide average nucleotide identity (ANI) compared to *B. pertussis* C734 was 82.6% and 98.8%, respectively. Recovered *B. holmesii* reads were limited to loci of high sequence homology and covered only 27.7% of the *B. pertussis* reference genome with 20X depth (Figure S2, Table S3). Significantly more *B. parapertussis* sequencing reads were recovered and mapped more evenly to cover 87% with 20X coverage (Figure S2, Table S3), consistent with the closer phylogenetic history of the species (10, 11, 34).

Taken together, the results from sequencing spiked specimens suggest a baseline limit of detection (LOD) for CIWGS at IS*481* Ct < 30 for 1x enrichment and IS*481* Ct < 35 for 2x enrichment. DNA extracts from residual pertussis-positive NP specimens with at least 10 ng total DNA should yield enough filtered sequencing read data sufficient for *B. pertussis* genomic characterization by SNPs or cgMLST/wgMLST (breadth 20x > 95%). Although genome fragments from closely related species are cross-reactive with the RNA bait library, filtered reads from *B. holmesii* or *B. parapertussis* will fail to meet minimum coverage thresholds or exceed expected SNP density, respectively.

### Surveillance specimens with matched isolates

Finally, CIWGS was evaluated with a set of 29 residual pertussis-positive NP specimens collected through the EPS program and from which cultured isolates were also available for independent sequence confirmation compared to a set of 28 unenriched NP specimens (TN). Diagnostic IS*481* Ct values exhibited by the selected specimens were consistent with those typically collected for laboratory confirmation during 2011-2019 (Figure S3). Filtered read recovery from the 29 surveillance specimens was similar to that observed from spiked negative specimens (Figure 3) and yielded sufficient data for genomic characterization with >80% breadth at 20X depth and >3,000 wgMLST allele calls (Table 1, Table S7, Figure S4).

**Table 1.** Comparison of matched specimen CIWGS and isolate sequencing.

| Specimen ID | IS481-Ct | DNA (ng/ul) | Num enrichments | Isolate ID | Isolate accession | CIWGS Breadth-20x | SNPs by Ref mapping | SNPs vs PacBio assembly | CIWGS Accession |
| --- | --- | --- | --- | --- | --- | --- | --- | --- | --- |
| EPS01 | 19.7 | 1.2 | 1 | J979 | SRR10002514 | 98.32 | 0 | NA | SRR27109067 |
| EPS02 | 22.7 | 2.5 | 1 | J984 | SRR10002466 | 95.30 | 0 | NA | SRR27109066 |
| EPS03 | 23.6 | 0.9 | 2 | J996 | SRR10002457 | 93.54 | 0 | NA | SRR27109065 |
| EPS04 | 22.3 | 6.0 | 1 | J968 | SRR9988730 | 95.42 | 0 | NA | SRR27109064 |
| EPS05 | 16.8 | 2.9 | 1 | J808 | SRR9131699 | 99.48 | 0 | NA | SRR27109063 |
| EPS06 | 25.3 | 2.5 | 2 | J836 | CP033293 | 87.44 | 0 | 0 | SRR27109061 |
| EPS07 | 14.5 | 0.1 | 1 | J841 | SRR9997167 | 99.83 | 0 | NA | SRR27109060 |
| EPS09 | 18.3 | 7.1 | 1 | J733 | CP046988 | 86.89 | 0 | 0 | SRR27109059 |
| EPS12 | 14.9 | 4.6 | 1 | J722 | SRR9131707 | 99.84 | 0 | NA | SRR27109058 |
| EPS14 | 16.3 | 0.1 | 1 | J726 | SRR9131600 | 97.70 | 0 | NA | SRR27109105 |
| EPS15 | 19.2 | 3.5 | 1 | J727 | SRR9131601 | 96.27 | 0 | NA | SRR27109104 |
| EPS17 | 21.7 | 3.2 | 1 | J788 | SRR9131623 | 94.93 | 0 | NA | SRR27109103 |
| EPS18 | 13.6 | 0.1 | 1 | J789 | SRR9131622 | 99.02 | 0 | NA | SRR27109102 |
| EPS20 | 12.0 | <0.1 | 1 | K056 | CP114533 | 99.70 | 0 | 0 | SRR27109101 |
| EPS22 | 21.6 | 7.6 | 1 | K712 | CP094044 | 94.21 | 0 | 0 | SRR27109100 |
| EPS24 | 27.8 | 11.8 | 1 | L021 | CP093981 | 99.81 | 0 | 0 | SRR27109098 |
| EPS28 | 18.0 | <0.1 | 1 | K921 | CP094014 | 99.39 | 0 | 0 | SRR27109097 |
| EPS29 | 14.3 | 12.0 | 1 | J866 | CP033281 | 99.92 | 0 | 0 | SRR27109096 |
| EPS31 | 18.2 | 10.8 | 1 | J816 | CP033295 | 99.81 | 0 | 0 | SRR27109095 |
| EPS32 | 17.2 | 2.1 | 1 | K439 | SRR10247384 | 99.77 | 0 | NA | SRR27109094 |
| EPS33 | 25.5 | 1.1 | 1 | K926 | CP094013 | 97.80 | 0 | 0 | SRR27109093 |
| EPS34 | 14.7 | <0.1 | 1 | K911 | CP094016 | 99.64 | 0 | 0 | SRR27109092 |
| EPS35 | 20.2 | 2.1 | 1 | K447 | SRR9718349 | 99.55 | 0 | NA | SRR27109091 |
| EPS36 | 23.0 | 24.2 | 1 | K124 | SRR10002473 | 94.70 | 0 | NA | SRR27109090 |
| EPS37 | 17.5 | 4.4 | 1 | K129 | CP114532 | 99.20 | 0 | 0 | SRR27109089 |
| EPS38 | 18.4 | <0.1 | 1 | K135 | CP114531 | 97.78 | 0 | 0 | SRR27109087 |
| EPS39 | 14.3 | 5.6 | 1 | K140 | CP114530 | 99.63 | 0 | 0 | SRR27109086 |
| EPS40 | 19.1 | 6.6 | 1 | K232 | CP114535 | 98.43 | 0 | 0 | SRR27109085 |
| EPS41 | 27.6 | 4.1 | 1 | K237 | CP114534 | 89.81 | 0 | 0 | SRR27109084 |

Sequence accuracy was assessed by comparing SNP calls between matched CIWGS and isolate sequencing reads relative to the reference assembly for C734. Detected SNPs were concordant, with all 29 pairs exhibiting 0 SNP differences (Table 1). Similarly, reference-quality assemblies were available for 15 isolates and additional SNP detection by directly mapping corresponding CIWGS reads also revealed 0 SNP differences (Table 1). The calculated percent agreement was 100% (TP = 29/29) and the overall percent agreement was 100% (TP = 29, TN = 28, FP = 0, FN = 0), according to calculations listed in the methods. To assess the consistency of CIWGS for genomic epidemiology, detected SNPs were used to reconstruct phylogenies from either the CIWGS or matched isolate data alone, as well as mixed together. Phylogenetic trees calculated from either CIWGS or isolate sequence data alone demonstrated concordant topologies, consistently clustering groups of putatively linked cases (Figure S5). Combined phylogenetic reconstruction of CIWGS and isolate sequence data accurately placed all matching pairs jointly in the resulting tree, also capturing potential linkages among the sampled cases predicted by either data type alone (Figure 4). However, core SNP alignments including CIWGS read data exhibited shorter sequence lengths, and thus effectively lower resolution, compared to those of isolate sequences alone (CIWGS: 134 bp, isolates: 173 bp, combined: 124 bp).

**Figure 4.**
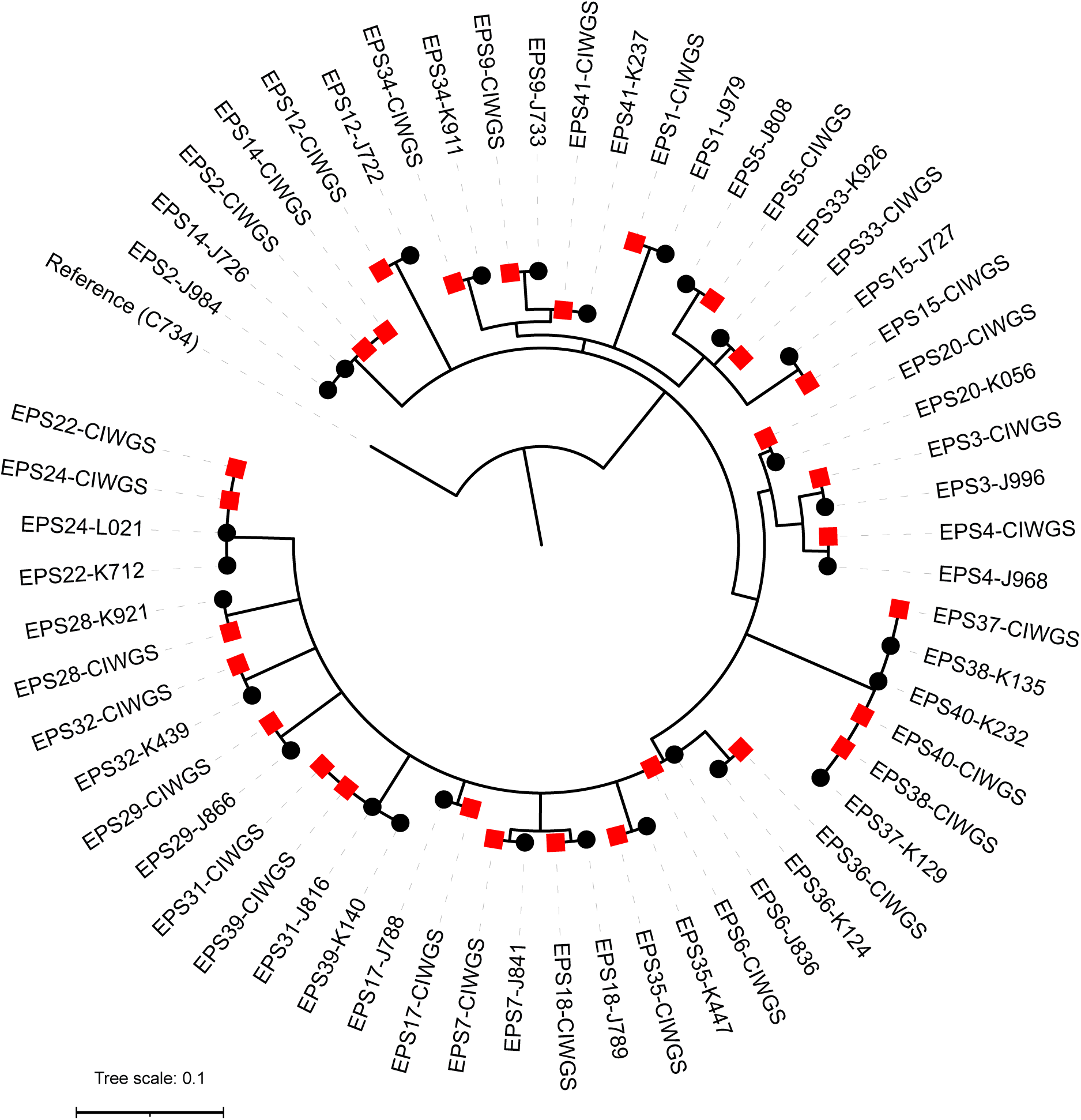
Phylogenetic reconstruction of matched CIWGS (red squares) and isolate (black circles) sequences from 29 surveillance specimens, reconstructed with 124 core, variable sites using maximum likelihood.

The matched data from these 29 surveillance cases were similarly compared by allele-based cluster detection using wgMLST. Allele profiles were concordant between CIWGS and isolate sequence data, with all pairs exhibiting 0 allele differences in a minimum spanning tree (Figure S6A, Table 1). However, clustering isolate sequences alone revealed additional segregation among the cases, concordant with the SNP phylogeny, whereas a tree of only CIWGS data closely resembled the combined tree (Figure S6B and S6C). As with the SNP alignments, wgMLST allele profile comparisons with CIWGS read data featured fewer available polymorphic loci for analyses (CIWGS: 49 loci, isolates: 99 loci, combined: 37 loci). Quality metrics indicated that CIWGS yielded poor *de novo* assemblies that were more fragmented and contained more ambiguous bases than those from matched isolate sequences (Figure S7A-D, Table S7), which negatively impact allele calling in BioNumerics.

Consensus allele calling rates from CIWGS data were often lower than from their matched isolate sequences (Figure S7E and S7F) and were not limited to specific scheme loci, suggesting that CIWGS data may yield lower resolution for wgMLST due to poor assembly performance.

### Detecting resistance mutations

Detection of antimicrobial resistance determinants by CIWGS was tested using two residual pertussis-positive NP specimens (PPHF 1999 and 3123) which putatively contained macrolide resistant *B. pertussis* (MRBP) as determined by PCR targeting mutant 23S rRNA alleles (35). Indeed, alignment of filtered CIWGS reads to 23S allele references confirmed a homozygous A2047G mutation of 23S in PPHF1999 and heterozygous A2047G mutation in PPHF3123 (Table S8). Further, filtered CIWGS reads from PPHF1999 also detected *fhaB3*, an allele primarily associated with MRBP isolated in east Asia (13).

## Discussion

We developed a target enrichment approach to CIWGS of *B. pertussis* directly from residual specimens used for positive clinical diagnosis. Advances in genomics and bioinformatics have modernized the study of microbial pathogens through reproducible, high-resolution strain characterization for molecular typing and epidemiology (9). However, concurrent widespread application of CIDTs like multiplex PCR or multi-pathogen panels have led to declines in available cultured isolates, often necessary inputs for WGS, from positive cases of bacterial pathogens like *B. pertussis*. The results presented here demonstrate the feasibility of CIWGS for accurate recovery of *B. pertussis* genomes from residual positive specimens that can be collected through an existing enhanced surveillance program. Application of these methods has the potential to strengthen surveillance, study the role of *B. pertussis* in disease resurgence, and monitor possible clinical impacts on vaccine-driven evolution by capturing additional cases with genomic characterization.

Although increased use of CIDTs has improved pertussis diagnosis and surveillance (14, 17), these gains have come at the cost of reduced availability of cultured *B. pertussis* isolates for laboratory characterization, including WGS. Increasing applications of clinical metagenomics may seem like an attractive alternative, capable of performing both detection and characterization in a single assay, but high levels of host patient contamination in NP specimens make pathogen genome recovery from shotgun sequencing challenging. The wide application of CIDTs, as well as variability among approved assay protocols (36, 37), currently used for pertussis diagnostics among US health laboratories makes replacement with metagenomic detection unrealistic.

The design approach implemented here intentionally focused on residual specimen material, which would often otherwise be discarded following diagnostic testing, to avoid impacting existing, validated protocols. To maximize the utility of available positive specimens during method development, additional surrogate specimens, prepared by spiking mock and negative specimens, were needed to optimize and evaluate the CIWGS workflow. Furthermore, focus on positive specimens both separates genomic characterization from variable case definitions and minimizes errors near the limit of detection. Previous evaluation of selective lysis demonstrated modest improvement in *B. pertussis* genome recovery from positive specimens (23) and similar results were obtained in preliminary testing prior to the development described here (data not shown). Instead, target enrichment takes advantage of the limited genetic diversity observed in *B. pertussis*, which often confounds conventional molecular assays (38), to capture conserved genome content. The required hybridization between RNA and single-stranded DNA depends on a minimal degree of sequence homology and the prepared RNA bait library must cover the expected level of divergence and content within the species, otherwise informative tracts of the pathogen genome may fail to be amplified for sequencing. Fortunately, the genome of *B. pertussis* remains highly conserved (10, 33, 39) and exhibits only slow genome reduction without any apparent gain of accessory genes (34, 40). The validation results presented here demonstrated that most of the *B. pertussis* genome, measured as coverage breadth at 20X depth, can be recovered by target enrichment with a bait library composed of DNA from only 10 diverse reference isolates. Other microbial pathogens with varied, open pangenomes may require careful reference selection or downstream analyses deliberately limited to conserved, core gene content.

CIWGS using target enrichment can produce data suitable for standard SNP or cgMLST/wgMLST analyses combined with existing *B. pertussis* isolate sequence data for both surveillance and research. However, evidence of intra-patient sequence diversity (33) may require subtle modification of downstream bioinformatics applications optimized for pure, cultured isolates. Adjusting minimum frequency thresholds to only capture predominant SNPs, or simply masking heterogenous sites altogether, may help but would simultaneously reduce overall resolution. While the results presented here illustrate concordant phylogenetic clustering with sequence data from matched isolates, analyses including CIWGS data did feature fewer variable sites or alleles for comparison. The resolution of wgMLST appeared to suffer most, which could be attributed to poorer *de novo* assembly of CIWGS data. Whether that makes CIWGS data better suited for genomic surveillance rather than genomic epidemiology remains to be seen, but the threshold for accurate cluster delineation may be as low as ∼2 alleles given the paucity of sequence diversity observed in *B. pertussis* (33, 39).

As expected, the RNA bait libraries also hybridized to and enriched highly conserved bacterial sequences (e.g., rRNA genes) as well as genes from closely related species of the genus *Bordetella*, consistent with current taxonomy (10, 34). Neither source of cross-reactive sequences diminishes the utility of the CIWGS or downstream characterization when using pertussis positive specimens as inputs, particularly when using diagnostic assays capable of accurately distinguishing *B. pertussis* from *B. holmesii* or *B. parapertussis* (37, 41).

Regardless, misidentification or even coinfection will be readily evident as clear aberrations from the expected low sequence diversity of true *B. pertussis* or detected using available metagenomic read classifiers (e.g., kraken2, mash).

While this novel approach may facilitate better understanding the epidemiology of pertussis and informing effecting prevention strategies, it is not without challenges. The optimized target enrichment laboratory workflow described here results in higher costs per sample compared to typical sequencing protocols for cultured isolates due to additional reagents, staff time, and equipment requirements. Minimal staff training is expected for laboratorians engaged in routine WGS on Illumina instruments. Rather, the most significant contributor to increased cost per sample is due to reduced multiplexing as CIWGS libraries require additional data output to achieve sufficient pathogen genome recovery, even after target enrichment. This cost discrepancy is expected to diminish as both sequencing costs and instrument output continue to improve.

Implementation faces other practical barriers such as timely access to residual diagnostic specimens, particularly from large commercial testing laboratories, and the variety of transport media or buffer solutions, nucleic acid extraction methods, and real-time PCR diagnostic assays currently used throughout the US public health testing network. It is also important to underscore the critical need for maintaining culture capacity at public health testing laboratories for *B. pertussis* antimicrobial resistance testing, monitoring acellular vaccine immunogen production, evaluating chromosome structural rearrangement (12, 42), and conducting animal challenge studies.

The widespread use of CIDTs and decline of culture threatens genomic surveillance of microbial pathogens, which in turn may undermine public health responses and control of infectious diseases. However, the prospect of genomic and phylogenetic analyses with larger datasets benefiting from the additional cases captured by CIWGS opens the door to addressing new questions about pertussis resurgence and the ecology and evolution of its etiologic agent *B. pertussis*.

## Materials and Methods

### Specimen and isolate sources

The Centers for Disease Control and Prevention’s (CDC) collection includes US *B. pertussis* isolates collected through surveillance and during outbreaks. Participating sites in the EPS program routinely submit both cultured isolates and residual NP specimens from pertussis-positive cases to CDC for confirmation and characterization, including whole-genome sequencing (17). Twenty-nine surveillance specimens for CIWGS validation were selected that met diagnostic input criteria and had matched isolates for comparison. Description of selected specimens are included in Table S7.

### Spiked specimen preparation

Mock clinical samples were prepared by mixing commercial, purified *Homo sapiens* DNA (Promega, Madison, WI) with a microbial community standard (Zymo Research, Irvine, CA) and then spiking purified *B. pertussis* isolate DNA from either E476 (CP010964) or K222 (CP114536) at varied ratios by mass (range = 0.000001 – 0.01) in duplicate. Separately, residual nasopharyngeal specimens, which were first confirmed negative for pertussis by diagnostic PCR (41), were pooled and 200 ul aliquots were spiked with a serial dilution of *B. pertussis* J865 (CP033408) cells (range = 0.1 – 10000) in triplicate.

### RNA bait library construction

A selection of 10 *B. pertussis* isolates, including two vaccine references and eight US predominant strains (Table S1), were used to generate a shotgun DNA template library for subsequent transcription to whole-genome RNA baits. Equal mass (1 ug) of genomic DNA from each of these 10 isolates was pooled, sheared, and then prepared into a whole-genome shotgun library using the NEB Ultra Library Prep kit (New England Biolabs; Ipswich, MA). Instead of the standard Illumina library prep adaptor, custom designed Y-type adaptors were ligated to both ends of the DNA fragments (Adaptor sequences: 5’— CGCTCAGCGGCCGCAGCACTTGAGAGAGAGAGATxT—3’, 5’-pATCTCTCTCTCTCAACCTCCTCCTCCGTTGTTG---3’).

After cleanup and size selection (300-500 bp) using AMPure XP beads (Beckman Coulter, Indianapolis, IN), the ligation products were amplified by 8 cycles of PCR on an S1000 thermocycler (Bio-Rad, Hercules, CA) using Q5 High-Fidelity PCR master mix with HF buffer (NEB Ipswich, MA) and PCR primers containing the T7 promoter sequence (Forward: 5’-GGATTCTAATACGACTCACTATACGCTCAGCGGCCGCAGCAC-3’, Reverse: 5’-GGATTCTAATACGACTCACTATACAACAACGGAGGAGGAGG-3’) (Adaptor and primers were synthesized in the CDC biotech core facility, Atlanta, GA). Initial denaturation was performed for 30 s at 98°C, followed by cycles for 10 s at 98°C, 30 s at 55°C, and 60 s at 68°C. The MEGAscript™ T7 Transcription Kit plus Bio-11-UTP (Invitrogen, Waltham MA) were used to produce the biotinylated RNA baits according to the manufacturer. The resulting RNA library was purified by using RNAClean XP beads (Beckman Coulter, Indianapolis, IN). The final quality and quantity were checked using a 2200 TapeStation with RNA Screen tapes (Agilent, Santa Clara, CA).

### DNA extraction, library preparation, bait hybridization, and sequencing

Isolates were cultured on Regan-Lowe agar without cephalexin for 72 h at 37 °C. Genomic DNA extraction from isolates was performed using the Gentra Puregene Yeast/Bacteria Kit (Qiagen; Valencia, CA) with slight modification (43). Briefly, two aliquots of approximately 1 x 10^9^ bacterial cells were harvested and resuspended in 500 uL of 0.85% sterile saline and then pelleted by centrifugation for 1 min at 16,000 x g.

Recovered genomic DNA was resuspended in 100 uL of DNA Hydration Solution. Aliquots were quantified using a Nanodrop 2000 (Thermo Fisher Scientific Inc.; Wilmington, DE). Genomic DNA extraction from clinical or mock specimens using the MagNA Pure platform (Roche Applied Science, Indianapolis, IN) following the method descripted in Burgos-Rivera *et al.* 2015 (36). Indexed Illumina sequencing libraries were constructed using the NEB Ultra Library Prep kit (New England Biolabs; Ipswich, MA).

Hybridization selection for CIWGS was conducted at 67.5°C for 24 h by incubating 0.5-1 μg of barcoded DNA libraries, 1 ug human DNA (Promega, Madison, WI), 5 ug sheared salmon sperm DNA (Promega, Madison, WI), 500 ng of RNA bait, and 25 ul Amersham™ Rapid-hyb Buffer (GE Healthcare) in a final volume of 50 μl using thermocycler (28, 32). After hybridization, captured DNA libraries were pulled down using Streptavidin Magnetic Beads (NEB). Beads were washed once at room temperature for 15 minutes with 0.5 ml 1 × SSC/0.1% SDS, followed by three 10-minute washes at 67.5°C with 0.5 ml pre-warmed 0.1 × SSC/ 0.1% SDS, resuspending the beads once at each washing step. Hybridized DNA library fragments were eluted with 50 μl 0.1 M NaOH. After 10 minutes at room temperature, the beads were pulled down, the supernatant transferred to a tube containing 70 μl of 1 M Tris-HCl, pH 7.5, and the neutralized DNA desalted and concentrated by using AMPure XP beads cleanup step. An additional PCR amplification was conducted using the corresponding barcode primer and universal primer with no more than 14 cycles. The final enrichened DNA libraries were sequenced using a MiSeq with the reagent v3, 500 cycle kit (Illumina; San Diego, CA).

### Bioinformatic read filtering

CIWGS reads were filtered to remove human sequence contamination by subtractive mapping to reference assembly GRCh38 using bowtie2 (v2.2.9), samtools (v1.3.1), and BEDTools (v2.26.0). In a similar manner, reads containing *B. pertussis* sequences were subsequently captured by mapping to reference C734 (CP013078). Summary statistics of target enrichment were calculated from the mapping output files.

### Sequence characterization and phylogenetics

The sequencing accuracy of filtered CIWGS reads relative to matched isolates was determined by mapping and SNP calling with snippy (v4.3.8) (https://github.com/tseemann/snippy) using the reference C734 (CP013078), and reference-quality assembly of isolate genomes if available, while masking all known IS elements. Maximum likelihood phylogenetic reconstructions were estimated from separate core SNP alignments of CIWGS and isolate data each, as well as combined, using RAxML (v8.1.16)(44). Tree visualization and annotation was performed with iToL (v6)(45). Tree structures calculated from CIWGS and isolate data individually were compared using phylo.io (46). Allele-typing by whole-genome MLST was performed with both CIWGS and isolate sequence data using BioNumerics (v7.6.3) as described previously (33).

Average Nucleotide Identity (ANI) between *B. pertussis* C734 and *B. parapertussis* J859 (CP043061) or *B. holmesii* C690 (CP020653) was calculated using the enveomics collection (46). Sequence similarly of all annotated protein coding genes in *B. parapertussis* J859 and *B. holmesii* C690 was evaluated by BLASTNn alignment (-qcov_hsp_perc 50) to the *B. pertussis* C734 reference.

### Performance characteristics

<u>Method accuracy was evaluated by comparing</u> read data from spiked specimens, both mock mixtures and pooled pertussis-negative NP aspirates, to previous reference-quality genome assemblies from the added *B. pertussis* isolate. True positive (TP) was defined as CIWGS data yielding >=85% coverage breadth at 20x depth and <= 2 SNPs relative to the reference genome, based on reported diversity among replicate *B. pertussis* isolates recovered during diagnostic culture (33). True negative (TN) was defined as either pooled, unspiked pertussis-negative NP aspirates or unenriched pertussis-positive surveillance specimens. Results were used in validation calculations:

Percent agreement = TP with positive result / TP total x 100 Overall percent agreement = TP+TN / TP+TN+FP+FN x 100

## Data availability

CIWGS shotgun sequencing reads without human sequence contamination are available from the NCBI Sequence Read Archive under BioProject accession number PRJNA922069 and isolate sequences are available under PRNJA279196.

## Code availability

Source code for scripts to further filter CIWGS sequencing read data are available at https://github.com/CDCgov/bpertussis-ciwgs/.

## Acknowledgements

We thank Pam Cassiday, Tami Skoff, and Matthew Cole (CDC) and The Enhanced Pertussis Surveillance/Emerging Infections Program.

This work was made possible through support from CDC’s Advanced Molecular Detection (AMD) program.

This activity was reviewed by CDC and was conducted consistent with applicable federal law and CDC policy.

The findings and conclusions in this report are those of the authors and do not necessarily represent the official position of the Centers for Disease Control and Prevention. Use of trade names and commercial sources is for identification only and does not imply endorsement by the Centers for Disease Control and Prevention, the Public Health Service, or the U.S. Department of Health and Human Services.

## Supplementary Figures and Tables

Figure S1. Optimization of hybridization temperature (A) and DNA library concentration (B) using mock mixtures spiked with *B. pertussis* isolate DNA.

Figure S2. Coverage mapping of *B. holmesii* C690 and *B. parapertussis* J859 vs *B. pertussis* C734 reference genome assembly; (A) *B. holmesii* C690 spike in recovery depth, (B) ANI of all C690 annotated genes vs C734 assembly, (C) *B. parapertussis* J859 spike in recovery depth, (D) ANI of all J859 annotated genes vs C734 assembly. Red line indicates genome-wide ANI.

Figure S3. Frequency distribution of IS*481* Ct values from typical EPS specimens received at CDC (black) compared to the 29 specimens analyzed here to evaluate CIWGS (grey).

Figure S4. Summary of *B. pertussis* recovery from 29 surveillance specimens [A] breadth-20x [B] wgMLST allele calls

Figure S5. Phylogenetic tree comparison between CIWGS (left) and isolate (right) data from the same 29 surveillance cases. Maximum-likelihood trees are constructed from 134 bp and 173 bp core, variable sites for CIWGS and isolate data, respectively.

Figure S6. Minimum-spanning trees of 29 surveillance specimens determined from wgMLST profiles of (A) CIWGS and isolates together, (B) isolates alone, or (C) CIWGS alone. CIWGS data are indicated in red, isolate data are indicated in grey. Circle diameter indicates cluster size. Labels on connecting edges indicate number of allele differences.

Figure S7. Quality metrics for *de novo* assembly and wgMLST allele calling of matched CIWGS (black) and isolate sequence data (red) from 29 surveillance specimens using BioNumerics. A full list of metrics is provided in Table S7.

Table S1. Genetic characteristics of *B. pertussis* strains used for RNA bait library preparation.

Table S2. Background microbial composition of pooled pertussis-negative NP aspirates by metagenomic read classification.

Table S3. Bordetella read content in pooled pertussis-negative specimens after removal of human sequence contamination compared to selected spiked and unenriched clinical specimens. Ratios, wgMLST stats, SNP calls.

Table S4. Sequencing read content of unenriched residual NP specimens.

Table S5. Comparison of parallel sequencing with and without enrichment.

Table S6. Comparison of parallel sequencing with 1x or 2x enrichment.

Table S7. Detailed recovery, *de novo* assembly, and wgMLST metrics for the 29 surveillance specimens and their matched isolates.

Table S8. CIWGS output from surveillance specimens with detected macrolide resistance mutations.

